# Comprehensive screening of a light-inducible split Cre recombinase with domain insertion profiling

**DOI:** 10.1101/2023.05.26.542511

**Authors:** Nathan Tague, Virgile Andreani, Yunfan Fan, Winston Timp, Mary J. Dunlop

## Abstract

Splitting proteins with light- or chemically-inducible dimers provides a mechanism for post-translational control of protein function. However, current methods for engineering stimulus-responsive split proteins often require significant protein engineering expertise and laborious screening of individual constructs. To address this challenge, we use a pooled library approach that enables rapid generation and screening of nearly all possible split protein constructs in parallel, where results can be read out using sequencing. We perform our method on Cre recombinase with optogenetic dimers as a proof of concept, resulting in comprehensive data on split sites throughout the protein. To improve accuracy in predicting split protein behavior, we develop a Bayesian computational approach to contextualize errors inherent to experimental procedures. Overall, our method provides a streamlined approach for achieving inducible post-translational control of a protein of interest.

## Introduction

The ability to control protein function is important for gaining insight into native biological systems and also engineering novel synthetic functions. Chemical inducers enable dose-responsive regulation of protein activity in response to small molecules, thus offering a versatile means of control. Optogenetic control offers an additional level of functionality since light can provide a reversible and precise input to enable temporal control over protein function. Both chemosensory and optogenetic approaches have been widely used in diverse fields ranging from biological discovery to engineering.^1–10^

Signal response times are particularly important for optogenetic protein tools because primary use cases center around controlling temporal changes.^11–15^ Because of this, post-translational mechanisms offer an attractive option for synthetic tools, because they avoid time delays associated with transcription and translation. Several studies have achieved post-translational control by splitting proteins with stimulus-responsive dimers.^2,7,16^ In the case of post-translational optogenetic control, light-responsive dimers are commonly used.^17–20^ In this approach, proteins are translated in two separate halves that are inactive. Each half is fused with a light-responsive dimer, which come together and dimerize in response to light exposure. For example, Nihongaki *et al*. introduced a Cas9 split with optogenetic dimers that can perform RNA-guided gene edits or gene repression in response to light.^10^ The split site where the optogenetic domains are incorporated within the protein of interest determines if the design is light-responsive, leaky, or non-functional. Deciding where to split the protein is often challenging, necessitating laborious screening of many split sites chosen based on structural information to find a functional construct.^7,10,21^ Computation approaches exist to predict functional split sites, which can assist in choosing candidates for testing.^22^ However, these models are not always accurate and do not take into account the dimer domains used, which can influence split site functionality.^21^

In principle, the challenge of identifying where to insert optogenetic domains is more tractable and limited in scope in comparison to other protein engineering endeavors, such as mutagenesis screens or directed evolution studies. The number of constructs to test is on the order of the number of amino acids in the protein of interest (hundreds to thousands). Consequently, we reasoned that the challenge of identifying functional split sites could be addressed efficiently with a parallelized approach by screening all possible split sites in a pooled library (Fig. 1).

**Figure 1.**
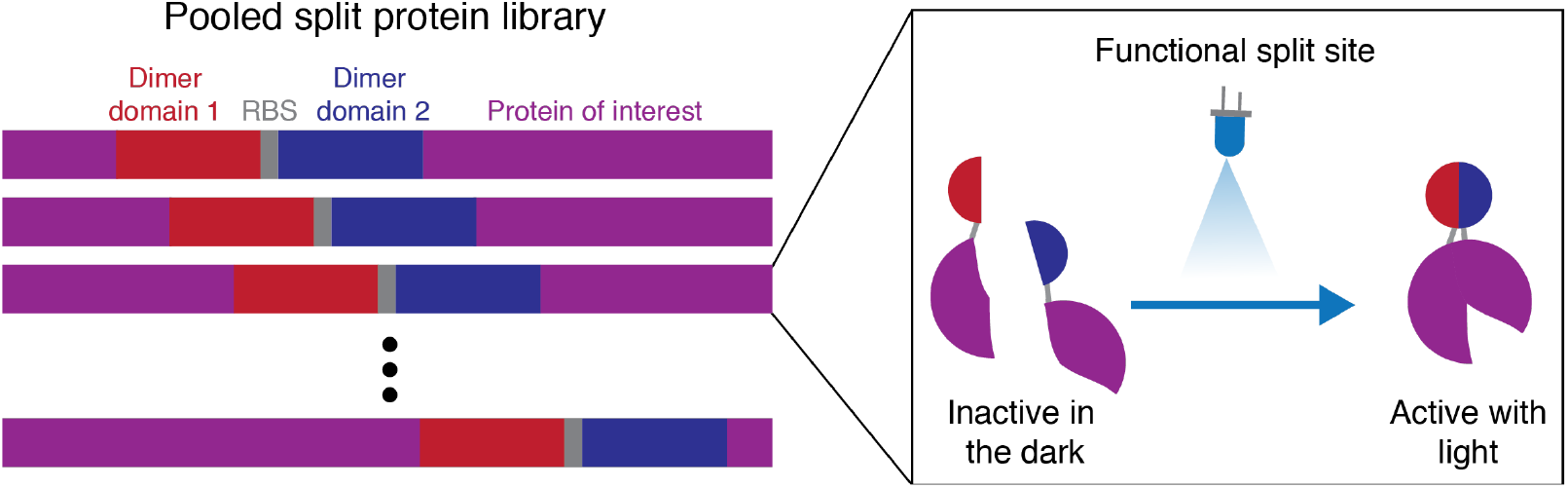
Schematic representation of the split protein library. A protein of interest is split at all potential amino acid sites and screened in parallel to identify functional split sites that are responsive to a specific stimulus. Here, the example of light-inducible dimers is used to illustrate how a split protein can be rendered light-responsive.

To simplify the library construction process, we used a pooled approach to generate variants with proteins split at all possible sites. Mahdavi *et al*. demonstrated that split libraries can be generated and screened with transposon mutagenesis.^23^ However, their work identifies split proteins that spontaneously combine, which is valuable in certain contexts, but not ideal for post-translation control of protein function. In order to generate libraries capable of input-responsiveness, domains also need to be inserted into the protein of interest with minimal scarring to facilitate functional translation and folding. Nadler *et al*. created a library generation method called domain insertion profiling combined with sequencing, or DIP-seq, that utilizes a MuA transposase to insert a modified transposon randomly within a sequence of interest (Fig. 2a).^24^ The transposon contains a chloramphenicol resistance (Cm^R^) gene flanked by BsaI sites, a type IIS restriction enzyme site that allows the Cm^R^ sequence to be swapped out for a desired insertion sequence at a later stage. This cloning scheme generates a library of constructs where a domain is inserted comprehensively into a plasmid to create a transposon insertion library. To isolate insertions that are within the gene encoding the protein of interest, BsmBI restriction enzyme sites flanking the region are used to excise the sequence and clone it into an expression vector using a golden gate reaction^25^ to create an expression library. The transposon fragment within these plasmids is then replaced with the domain that will be incorporated into the protein of interest. In their study, Nadler *et al*. inserted a fluorescent protein domain into a protein that binds trehalose to create a fluorescent biosensor.^24^ We reasoned that the DIP-seq technique could be modified to create split protein libraries by instead inserting two dimerization domains with an internal ribosome binding sequence (RBS), such that the resulting constructs are expressed as two proteins (Fig. 2b).

**Figure 2.**
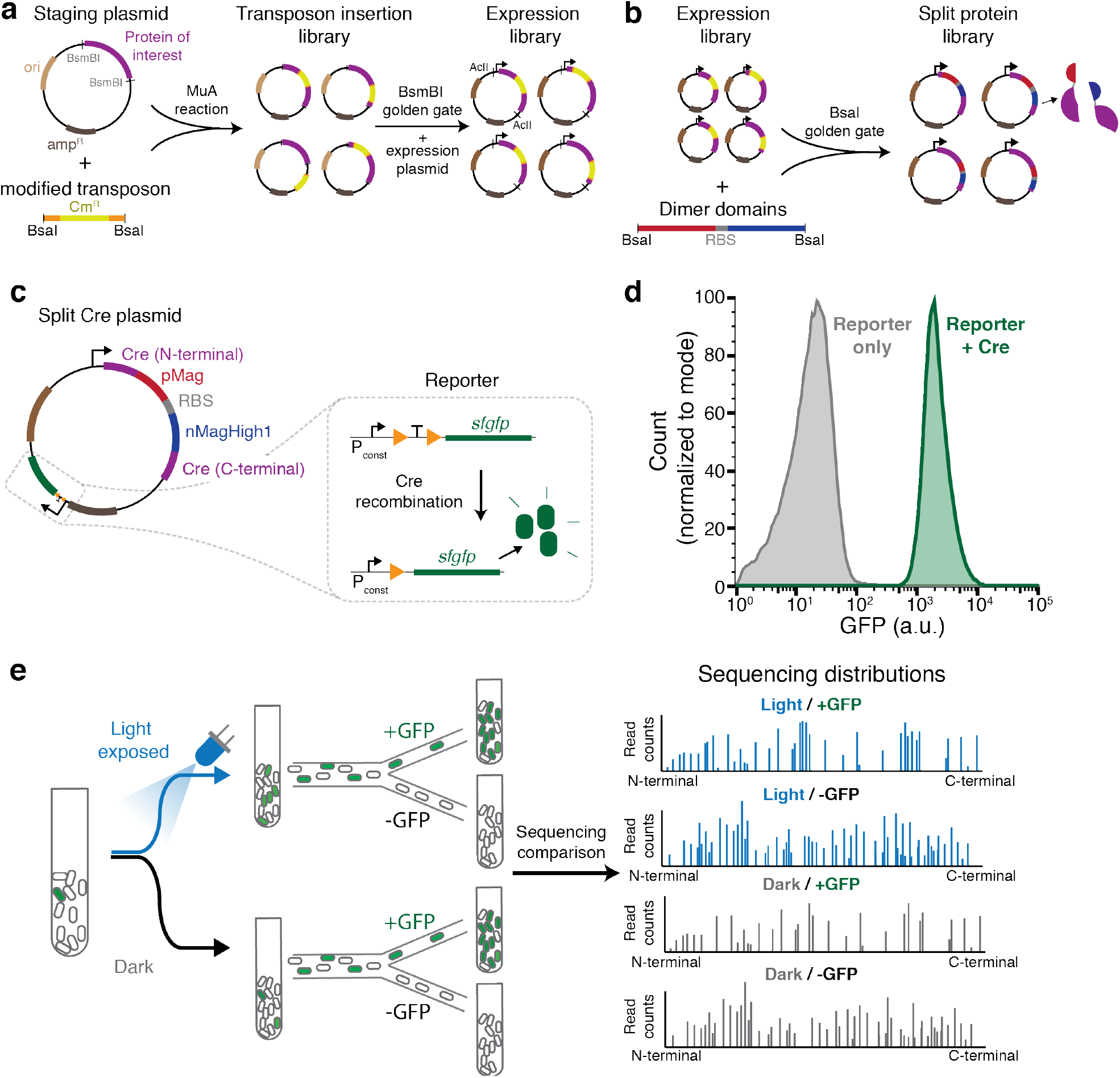
Generating and screening split protein libraries. **(a)** Method for generating expression libraries with a modified transposon with flanking BsaI restriction enzyme sites. MuA transposase reactions insert the modified transposon containing the chloramphenicol resistance gene (Cm^R^) randomly into the staging vector containing the gene encoding a protein of interest to create a transposon insertion library. The BsmBI sites are then used for a digestion and gel extraction to isolate the sequence encoding the protein of interest containing a transposon insertion. This fragment is used in a golden gate assembly into the expression vector to generate the expression library. AclI sites are used to digest the library prior to sequencing. **(b)** The modified transposon in the expression library is then replaced by dimerization domains with an internal RBS through a BsaI golden gate reaction to generate a split protein library. **(c)** Each library variant has Cre split by the light-inducible dimers pMag and nMagHigh1. The expression vector for the split Cre library contains a recombinase activity reporter. Cre recombinase excises a transcriptional terminator placed between a constitutive promoter, P_const_, and the gene encoding sfGFP, resulting in an increase in fluorescence when Cre is functional. **(d)** Fluorescent histograms of the Cre expression reporter with and without Cre. **(e)** Overview of the experimental workflow used to identify functional split sites within the split Cre library. Libraries are kept in the dark or exposed to 465 nm blue light and sorted based on fluorescence. Sorted libraries are sequenced and compared to distinguish activity of each split site. Library distributions from light exposed and dark populations are compared to uncover individual split site behavior.

Here, we demonstrate a method of creating and screening split proteins in pooled libraries. As a proof of concept, we profiled split sites for Cre recombinase, a widely-used protein in genetics and synthetic biology that enables site-specific gene recombination, inversion, or excision.^26,27^ We split Cre and inserted light-responsive dimer domains, called ‘magnets’, and comprehensively profiled for activity in response to blue light.^20^ We then used pooled sequencing to link split site library distributions to functional behavior of individual split proteins. To analyze the datasets output from our workflow, we designed a Bayesian inference approach to output predictions on the functionality of individual split sites. This approach is capable of managing several sources of error that arise experimentally, such as heterogeneous library distribution and spurious cell sorting events, and provides a probabilistic prediction about whether a given split site is likely to be a functional hit. We validated the approach by individually characterizing several split proteins that the model predicted to be hits, leaky variants that are not light-responsive, or non-functional results. In sum, we describe a method to create post-translational optogenetic control. This approach should be generalizable to other proteins or inducible dimer domains of interest.

## Results

### Design of parallel split Cre screening

To create a split Cre recombinase library, we applied the modified domain insertion approach to construct a staging plasmid containing the Cre coding sequence with a 17 amino acid N-terminal truncation as described in Jullien *et al*.^2^ (Tables S1-S3). We created a transposon insertion library by integrating the modified transposon randomly into a Cre staging plasmid using MuA transposase, as described above. Once the Cre-transposon fragment was cloned into an expression vector, the transposon sequence was replaced by an insertion fragment containing the optogenetic ‘magnet’ heterodimers (pMag and nMagHigh1) with an RBS between each dimer (Fig. 2c).^20^ The insertion fragment was designed such that, when inserted in-frame, glycine-serine linkers are incorporated between the Cre coding sequence and each magnet (Table S2-S3). A five-nucleotide sequence from Cre is duplicated during the transposon reaction and the insert fragment was designed to maintain an open reading frame for the N-terminal and C-terminal split Cre fragments (Fig. S1). Sequencing of the initial split Cre expression library confirmed that the Cre sequence was comprehensively split, with >85% (282 out of 326 codons) of total split possibilities present.

As a readout to measure recombination activity, we designed the expression vector to contain a fluorescent reporter (Fig. 2c). The reporter contains a transcriptional terminator flanked by *loxP* sites placed in between a constitutive promoter and the gene encoding superfolder GFP (*sfGFP*). Without active Cre, the terminator blocks transcription of *sfgfp*. Cre-mediated recombination excises the terminator resulting in a 129-fold increase in green fluorescence (Fig. 2d).

Using our expression library containing Cre recombinase split at the majority of amino acids, we next subjected the library to an experimental workflow devised to test the behavior of each variant of split Cre in parallel (Fig. 2e). Since the expression plasmid also contains a fluorescent reporter for Cre activity, the activity of a given split Cre variant can be linked to the sequence that generated the cutting. Using fluorescence-activated cell sorting (FACS), we sorted cells based on Cre activity and sequenced the plasmid populations to determine which split Cre variants were present. Specifically, we divided the pooled library into two equal parts and subjected one to blue light exposure while the other was kept in the dark. Using FACS, we separated cells that expressed GFP (denoted +GFP) and from those that did not (-GFP) for each light treatment condition. By treating the library with blue light or keeping it in the dark prior to sorting, we reasoned that we could compare populations to determine which split sites were light-responsive (Fig. 2e). We performed next generation sequencing on each of the four libraries (Light/+GFP, Light/-GFP, Dark/+GFP, Dark/-GFP) and filtered for reads that contained the interface between the Cre coding sequence and either pMag or nMagHigh1 to identify the split protein variants present in each library. Counts for these filtered reads were greater than 10^4^ for each of the four libraries.

### Experimental error and Bayesian modeling

We considered three possible phenotypes for each split Cre variant: a hit (active in the light but not in the dark), leaky (active even in the dark), or non-functional (inactive in the light and the dark) (Fig. S2). Our initial hypothesis was that by comparing the Light/+GFP population to the Dark/+GFP population, it would be straightforward to identify hits. However, initial attempts at analyzing these populations through simple measures, such as dividing the proportion of each split site in the Light population by the proportion in the Dark population to identify hits proved inaccurate. Simple analysis measures like this were prone to false positive hit predictions stemming from low coverage of certain sites due to low sequencing read counts associated with some split variants, with low or null read counts on either Light or Dark populations strongly affecting these ratiometric calculations. Therefore, we examined our experimental workflow for sources of error that could be causing inaccurate predictions and identified three major factors that can contribute to error. First, variants are represented in the naïve libraries prior to cell sorting with widely heterogeneous abundances, spanning several orders of magnitude in relative proportion (Fig. S3). Consequently, a substantial proportion of variants are present in low quantities because of uneven sampling of each variant. In an ideal scenario, this distribution would be uniform across the split sites. Second, the sorting process happens with a substantial error rate, which we estimate to be approximately 25%, and is likely to sort many variants into the wrong populations (i.e. a variant that is -GFP will be sorted into +GFP about 25% of the time). Third, the sequencing process draws sequencing reads randomly from each population distribution, which introduces intrinsic sampling noise.

To manage these experimental realities, we expanded our workflow and analysis in two directions. First, we used sequencing data not just from Light/+GFP and Dark/+GFP populations, but also Light/-GFP and Dark/-GFP populations. By incorporating these additional populations, we were able to utilize several more comparisons to double check if a conclusion from a single comparison was the result of an error (Fig. S4). For example, comparing the Light/-GFP to Dark/-GFP may also reveal hits through the absence of variants in the Light/-GFP population. Comparing Light/+GFP to Dark/-GFP can be used to reveal leaky variants. Second, we incorporated Bayesian modeling into our analysis, which assigns phenotypic probabilities to each split site based on sequencing read counts to predict performance of each split (Supplementary Text). Bayesian modeling provides a framework for incorporating all the data from the naïve and sorted populations to produce predictions that combine all the available information, instead of relying on empirical pairwise comparisons between populations. A custom Bayesian model additionally allowed us to explicitly describe the sources of uncertainty present in the system, including those that are intrinsic or result from inaccurate observations. Finally, the fact that the results are probabilistic reflects the degree of confidence that one is allowed to expect from a particular dataset: a split site represented in low abundance in the population will result in predictions that do not differ much from the uninformed hypothesis, while an abundant split site might be associated with a more confident prediction.

### Split Cre mapping and validation

Using our data analysis improvements, we were able to predict the performance of nearly all split sites in the library, resulting in a Cre ‘split map’ (Fig. 3a). In our Bayesian approach, each split site is designated with a probability for which it will behave as a hit, leaky, or non-functional variant. As an initial sanity check, we noted that our predictions aligned well with known split Cre data from literature. For example, the split at amino acid 42 has been validated as a light-responsive Cre construct with magnets.^21,28^ Additionally, a study from Weinberg *et al*. identified multiple functional split sites in the region between 210-260, which align with the general trends we observe in the split map.^7^ However, it is important to note that there are differences between the split constructs tested here and reports from the literature. First, a 5-nucleotide duplication caused during the transposon reaction adds two extra amino acids to the C-terminal compared to traditional split proteins (Fig. S1). Also, the expression level and organism likely influence split performance and are specific to each experimental set up.

**Figure 3.**
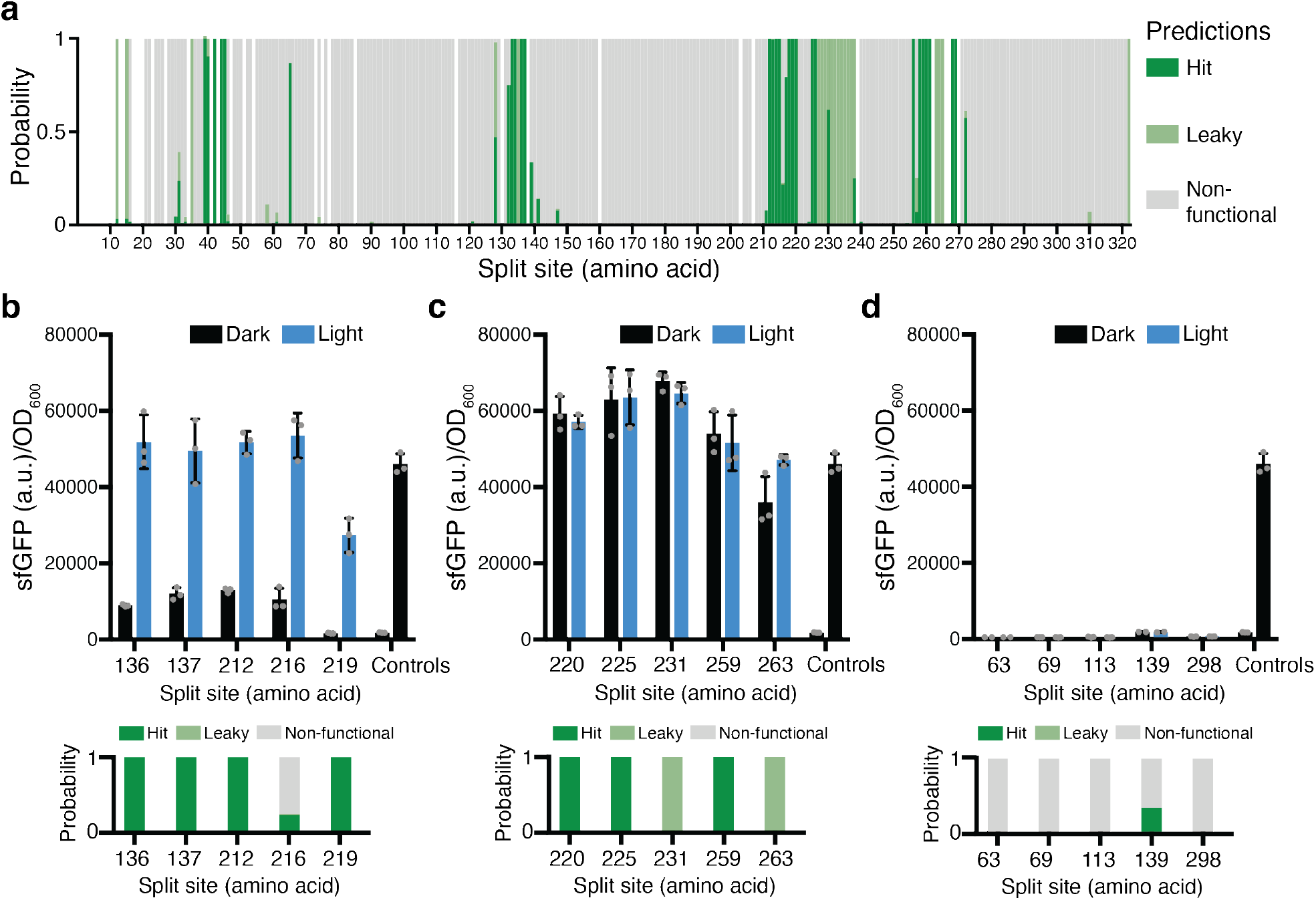
Cre recombinase split map predictions and validation. **(a)** Phenotypic predictions of split Cre at various amino acid split positions. Each split Cre in the library is assigned a probability of behaving as a hit (dark green), leaky (light green), or non-functional (gray) variant. Split sites that were missing from either naïve library are left white. **(b-d)** Validation of individual split Cre constructs. sfGFP expression levels with and without exposure to 465 nm blue light are compared to predictions from the library screen to identify true **(b)** hits, **(c)** leaky, or **(d)** non-functional split variants. Controls display sfGFP expression from the reporter with and without a functional, non-split Cre. Error bars show standard deviation around the mean (n = 3 or 2 biological replicates; individual data points are overlaid on the bar plot).

To assess the quality of the split map predictions, we constructed several individual split Cre variants to test light responsiveness. We built 15 different variants, focusing on predicted hits, but also including some predicted leaky variants, variants where the model prediction was uncertain, and some predicted non-functional variants. Of these 15 variants, we identified five split sites that were light-responsive (Fig. 3b). Four out of five of these hits—splits at amino acids 136, 137, 212, and 219—were predicted as such. One hit—amino acid 216—was predicted to behave as a non-functional split, however, the model estimated a 22% probability that this split would behave as a hit. An additional five constructs were leaky during individual validation experiments (Fig. 3c). In this case, two of five leaky splits were predicted accurately as leaky, while three were predicted to be hits. Therefore, the model is prone to false positives when predicting light responsiveness but performs well when predicting overall catalytic activity. In addition, we found constructs that behaved as non-functional variants (Fig. 3d). In this case, five out of five non-functional splits were predicted correctly in the Bayesian analysis.

During the validation process, we uncovered a degree of heterogeneity between colonies. Clonal replicates behaved similarly, as evidenced by good reproducibility between replicates (Fig. 3b-d). However, separate colonies of the variant with the split at amino acid 219, which is a hit, responded differently to light (Fig. S5). Clonal replicates are taken from the same colony after cloning and separated into different cultures during functional testing. Alternatively, separate colonies can be taken during the cloning process and used as separate replicates. In all cases, plasmids were sequenced to verify accurate construction. In the case of 219, one of four colonies displayed a leaky phenotype, while all others were light-responsive. The leaky phenotype may stem from the non-reversible nature of the reporter. If the reporter is activated early during the cloning process, either through an active Cre in the dark state or accidental ambient light exposure, then the +GFP signal will be maintained indefinitely. The heterogeneity in behavior for an individual split may contribute to the prediction errors we encountered when analyzing the sequencing dataset because a given split may behave as several phenotypes. This reality further motivates a Bayesian analysis approach that assigns probabilities instead of making a discrete phenotypic assignment.

Interestingly, areas of catalytic activity, whether leaky or a hit, appear in clusters within the Cre sequence (Fig. 3a). To gain structural understanding, we mapped catalytic activity to the structure of Cre recombinase (Fig. 4a). The functional clusters are primarily in loops between secondary structures, which may be less likely to disrupt protein function. Also, areas of the binding interface between monomers all appear as non-functional variants. In this case, steric hindrance from the dimerization domains may interfere with tetramer formation, rendering the construct non-functional. Notably, four of five clusters appear on one plane of the Cre structure relative to the bound DNA (Fig. 4b). In addition, we plotted catalytic activity of each split site against B-factor, which captures disorder in the crystal structure,^29^ however we did not observe a clear correlation with activity indicating it cannot be used to aid functional split site predictions (Fig. S6). Overall, these results demonstrate that our method not only reveals split Cre hits, but supplies holistic protein information for hit, leaky, and non-functional phenotypes that may be useful for informing further engineering of Cre or proteins with similar structure.

**Figure 4.**
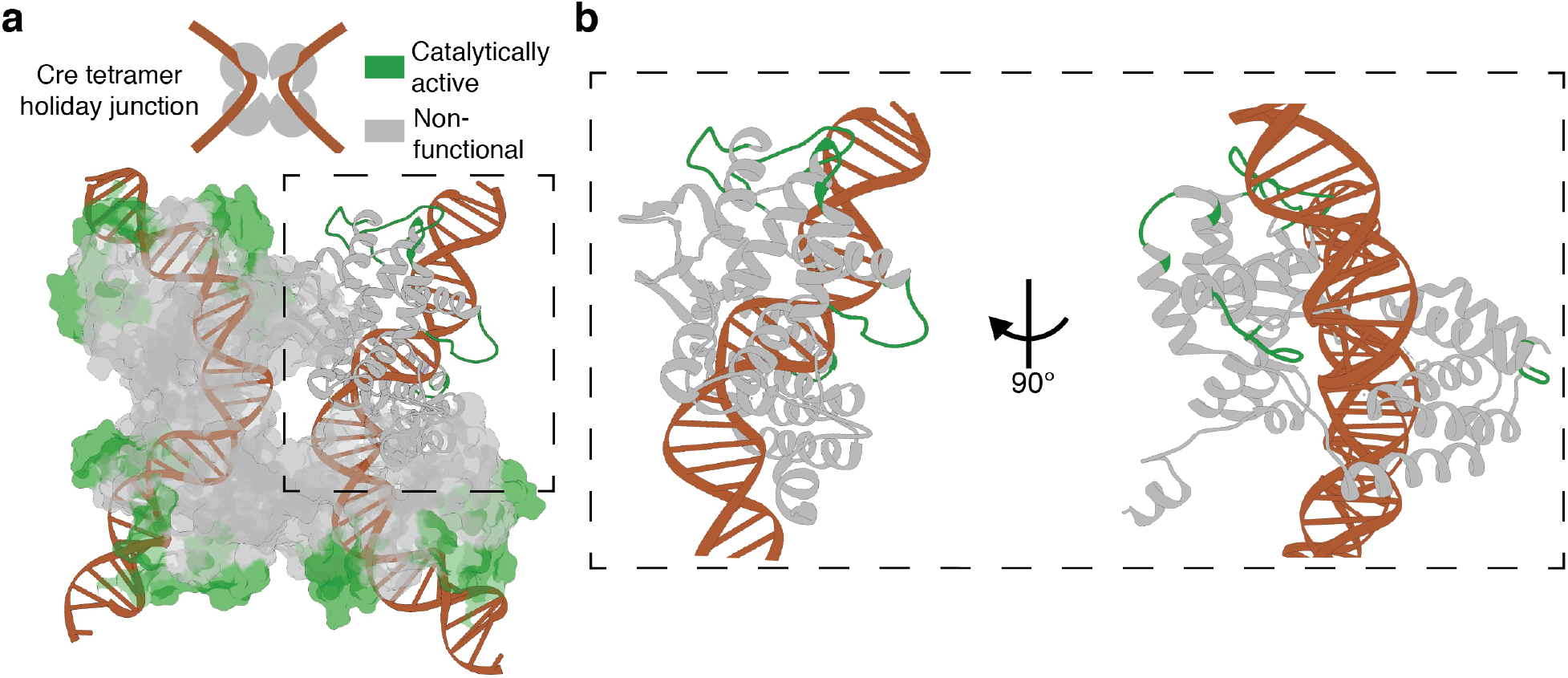
Split map predictions and protein structure. **(a)** Regions with several residues predicted to be catalytically active when split (hit or leaky, highlighted in green) are mapped to the structure of tetrameric Cre bound to DNA (PDB: 1NZB). Regions highlighted: aa39-45; aa132-137; aa212-220; aa225-238; aa256-269. aa, amino acid. **(b)** Zoomed in structure of a Cre monomer with catalytically active split sites highlighted in green.

## Discussion

Here, we demonstrate a method of creating and screening split protein libraries. With this approach, we provide a technique for incorporating post-translational optogenetic control into proteins without the need for detailed protein engineering expertise or uninformed screening of variants. Our approach can, in principle, be extended to other proteins of interest given cases where it is possible to screen or select for functional variants. Although we focus on light responsiveness, we envision that the experimental workflow and dimer system can be modified to create chemically responsive split proteins, as well.

In this work, we comprehensively screened Cre recombinase split with ‘magnet’ dimers as a proof of concept. In addition to matching trends from previous split Cre studies, we were able to identify several novel light-responsive split sites. We identified intrinsic sources of error in the experimental workflow that contribute to inaccurate predictions when using basic analysis methods. By enhancing our sampling and incorporating a Bayesian model of the experimental error, we make fairly accurate predictions despite these uncertainties in the dataset. We believe this approach to modelling experimental error could extend to other experimental workflows where non-uniform library coverage or incomplete readouts are inherent.

Comprehensively splitting a protein leads to greater structural insight compared to testing only a subset of variants predicted to function. As demonstrated with Cre, data for each amino acid split variant present in the library can be mapped to structure to identify trends. We found that loops distant from a dimer interface are optimal to avoid disruption of recombination activity. Extending our approach to several proteins using various dimerization systems may reveal generalizable rules that could inform split protein engineering strategies. Datasets generated by parallel approaches such as ours could also be used to inform computational methods to improve model accuracy.^22,30,31^

An interesting finding we encountered through validation of individual split constructs was the potential for a split to have heterogenous behaviors between colonies (Fig. S5). Importantly, our prediction model is capable of capturing this variation. It is possible that early expression variation during the cloning process may lead to varying degrees of reporter cutting that are propagated through the validation experiment. Performing the screen with a low-copy plasmid with a more tightly controlled promoter may overcome this issue in the future. Also, we chose the magnet variant nMagHigh1 which contains a kinetic mutation to increase dimer half-life.^20^ This may sensitize the library to accidental ambient light exposure, causing validation heterogeneity.

Our method has several limitations. A downside to creating domain insertion libraries with the MuA reaction is that the library distribution is highly heterogeneous, meaning each variant is represented unevenly within the library. Additionally, from a probabilistic standpoint, the majority of the variants produced are not in frame, which decreases sequencing depth. Consequently, our prediction accuracy suffered from false positive predictions of several hits that were found instead to be leaky during validation. Experimental modifications could improve phenotype prediction accuracy. For example, an alternative method for creating domain insertion libraries, called SPINE, uses an oligo library approach instead of transposons.^32^ Using SPINE for split protein libraries could produce a higher quality starting library, albeit at a higher cost of library creation.

Compared to an off-the-shelf statistical method, our custom Bayesian model is flexible and makes it possible to incorporate the precise assumptions or expert knowledge about the problem or the experimental setup. For example, we assumed that the two naïve distributions are distinct enough to warrant modelling them as separate, but modelling them as a unique distribution is possible, and would yield slightly different results. We also fixed values for the false positive and negative rates of the FACS process, but we could consider inferring values for these rates themselves, at a computational price related to a more difficult sampling.

In summary, this work demonstrates a method to identify functional split sites. Through parallel screening of many split proteins, it allows researchers to confer post-translational control that is agnostic to the protein of interest while requiring little protein engineering expertise and reducing the validation necessary to identify functional variants.

## Methods

### Strains and plasmids

We created transposon insertion libraries where the chloramphenicol resistance (Cm^R^) gene *cat* was inserted randomly in the staging plasmid as described in Nadler *et al*.^24^ using the following plasmids with some modifications: pUCKanR-MuA-BsaI (Addgene #79769), pATT-Dest (Addgene #79770), and pTKEI-Dest (Addgene #79784). The following modifications were made to the plasmids from Nadler *et al*. when creating insertion libraries: An mRFP1 sequence from pBbE5c-mRFP1 from the BglBrick library^33^ was added to pATT-Dest using golden gate assembly^25^ to create pATT-Dest-RFP (Table S1). This provided an easily distinguishable difference in insert size during gel extraction when excising Cre-transposon fragments from the pATT-Dest backbone. The Cre coding sequence was added to pATT-Dest-RFP to create the staging vector pATT-Dest-RFP-Cre (Table S1). The expression plasmid pTKEI-Dest was modified using golden gate assembly^25^ to place a Cre GFP reporter into the backbone to create pTKEI-Dest-loxP-sfGFP (Table S1).

The pMag-RBS-nMagHigh1 sequence was introduced on a separate pATT-Dest plasmid to create pATT-BsaI-pMag-nMagHigh1 that contains flanking BsaI sites, GS linkers, and the pMag-RBS-nMagHigh1 sequence (Table S1-S3). The magnet insert was amplified by PCR, followed by a PCR cleanup before being included in a golden gate reaction with the expression library to create the final split protein library. At all steps in the library generation process, colony forming unit (CFU) counts were performed on plates containing appropriate antibiotics to confirm that the number of transformants was >10^5^ to maintain coverage at least an order of magnitude above the number of possible insertions. A positive control plasmid containing non-split Cre was created by replacing the mRFP1 gene from the BglBrick arabinose inducible vector, pBbS8c-mRFP1, with the Cre coding sequence.^33^ Antibiotic concentrations used for plasmid maintenance were 30 μg/mL for kanamycin, 100 μg/mL for carbenicillin, and 25 μg/mL for chloramphenicol.

### Blue light stimulation

All light exposure experiments were carried out with a light plate apparatus (LPA)^34^ using 465 nm blue light. The split Cre library was cultured in Luria broth (LB) with appropriate antibiotics for plasmid maintenance at 37 °C with 200 rpm shaking and kept in the dark unless being exposed to blue light in the LPA. Overnight cultures of the split Cre library were diluted 1:500 and precultured in the dark for 2 hours. The library was then either kept in the dark or exposed to 100 μW/cm^2^ blue light for 2 hours followed by 3 additional hours in the dark before being diluted again 1:1000 for overnight growth prior to cell sorting. During validation experiments, single split variants were treated with the same culture and light exposure procedure. pBbS8c-Cre was co-transformed with pTKEI-Dest-loxP-sfGFP to serve as a positive control for Cre activity. Green fluorescence (excitation 480 nm, emission 510 nm) and optical density (OD at 600nm) readings were taken using a BioTek Synergy H1m plate reader after light induction or dark culturing and overnight growth, as described above.

### Cell sorting

Fluorescence-activated cell sorting was carried out on a Sony SH800S cell sorter using a 70 μm microfluidic chip. Singlet cells were gated based on GFP fluorescence of pTKEI-Dest-loxP-sfGFP without Cre introduced and pTKEI-Dest-loxP-sfGFP co-transformed with pBbS8c-Cre. Each culture, either kept in the dark or light exposed, was sorted for 200,000 +GFP cells on ultra-purity mode. A total of six libraries: +GFP, -GFP, and unsorted (naïve) cultures for Light or Dark conditions were miniprepped for sequencing.

### Sequencing and alignment

Prior to next generation sequencing, each library miniprep was digested with AclI and size-selected through gel extraction to only contain the portion of the plasmid with Cre and inserted magnets. Sequencing libraries were prepared using the Illumina DNA Prep kit, according to manufacturer’s specifications. Libraries were then sequenced on an Illumina MiSeq using 2×150 v2 chemistry. Sequencing reads were trimmed of adapters using Trimmomatic^35^ v0.39, and a custom Python script was used to identify Cre insertion loci. Out of frame insertions were filtered out and not included in downstream analysis. Splits were indexed (1-326) according to the amino acid sequence in Table S3.

### Bayesian modeling

The Bayesian approach uses a generative model to map various sources of uncertainty in the experimental process. Each split site can be either a hit (H), leaky (L), or non-functional (N). As a prior, for every site i from 1 to 326 we used:

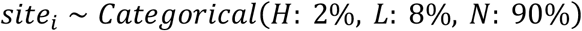

This reflects our expectation that most split sites will be non-functional and hits will be comparatively rare. We modeled the distribution of split sites in each naïve library (dark and light exposed, pre-sorting) as vectors of non-negative values summing to 1, distributed according to a Dirichlet distribution, which is in good agreement with the sequenced frequencies (Fig. S7):

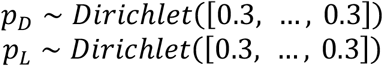

*p*_*D*_ is the dark naïve distribution and *p*_*L*_ is the light naïve distribution. The model accepts the six read count distributions (Light naïve, Dark naïve, Light/+GFP, Light/-GFP, Dark/+GFP, Dark/-GFP) and returns samples from the posterior distribution of the variables to estimate: the 326 *site*_*i*_, *p*_*D*_, and *p*_*L*_. We tally the samples for each *site*_*i*_ to compute hit, leaky, and non-functional probabilities. The Bayesian analysis was modeled and sampled by PyMC, a Python Bayesian inference library.^36^ An expanded explanation of our modeling approach is provided in the Supplementary Text.

## Data Availability

All sequence data are available in the Sequence Read Archive, under BioProject PRJNA940279.

## Code Availability

Code used for sequence analysis is available at https://github.com/timplab/tague_splitcre. Code used for Bayesian analysis is available at https://gitlab.com/dunloplab/split-cre-inference.

## Acknowledgments

We thank Dr. Luis Ortiz, Dr. Uros Kuzmanovic, and Dr. James Galagan for training and assistance with the FACS instrument; Michael Sheets for providing a Cre recombinase reporter which we modified for this study; and Dr. Dana Nadler for help troubleshooting transposon library creation experiments. This work was supported by NIH grant R01AI102922 and DOE grant DE-SC0019387.

## Notes

### Competing Interest Statement

The authors have declared no competing interest.

## References

(1) Feil, R.; Brocard, J.; Mascrez, B.; Lemeur, M.; Metzger, D.; Chambon, P. Ligand-Activated Site-Specific Recombination in Mice. Proceedings of the National Academy of Sciences of the United States of America 1996, 93 (20), 10887–10890. https://doi.org/10.1073/pnas.93.20.10887.

(2) Jullien, N.; Sampieri, F.; Enjalbert, A.; Herman, J. P. Regulation of Cre Recombinase by Ligand-Induced Complementation of Inactive Fragments. Nucleic acids research 2003, 31 (21). https://doi.org/10.1093/nar/gng131.

(3) Olson, E. J.; Tabor, J. J. Post-Translational Tools Expand the Scope of Synthetic Biology. Current Opinion in Chemical Biology 2012, 16 (3–4), 300–306. https://doi.org/10.1016/j.cbpa.2012.06.003.

(4) Davis, K. M.; Pattanayak, V.; Thompson, D. B.; Zuris, J. A.; Liu, D. R. Small Molecule-Triggered Cas9 Protein with Improved Genome-Editing Specificity. Nature Chemical Biology 2015, 11 (5), 316–318. https://doi.org/10.1038/nchembio.1793.

(5) Rose, J. C.; Stephany, J. J.; Valente, W. J.; Trevillian, B. M.; Dang, H. V.; Bielas, J. H.; Maly, D. J.; Fowler, D. M. Rapidly Inducible Cas9 and DSB-DdPCR to Probe Editing Kinetics. Nature Methods 2017, 14 (9), 891–896. https://doi.org/10.1038/nmeth.4368.

(6) Tague, E. P.; Dotson, H. L.; Tunney, S. N.; Sloas, D. C.; Ngo, J. T. Chemogenetic Control of Gene Expression and Cell Signaling with Antiviral Drugs. Nature Methods 2018, 15 (7), 519–522. https://doi.org/10.1038/s41592-018-0042-y.

(7) Weinberg, B. H.; Cho, J. H.; Agarwal, Y.; Pham, N. T. H.; Caraballo, L. D.; Walkosz, M.; Ortega, C.; Trexler, M.; Tague, N.; Law, B.; Benman, W. K. J.; Letendre, J.; Beal, J.; Wong, W. W. High-Performance Chemical- and Light-Inducible Recombinases in Mammalian Cells and Mice. Nature Communications 2019, 10 (1). https://doi.org/10.1038/s41467-019-12800-7.

(8) Dagliyan, O.; Tarnawski, M.; Chu, P. H.; Shirvanyants, D.; Schlichting, I.; Dokholyan, N. V.; Hahn, K. M. Engineering Extrinsic Disorder to Control Protein Activity in Living Cells. Science 2016, 354 (6318), 1441–1444. https://doi.org/10.1126/science.aah3404.

(9) Guntas, G.; Hallett, R. A.; Zimmerman, S. P.; Williams, T.; Yumerefendi, H.; Bear, J. E.; Kuhlman, B. Engineering an Improved Light-Induced Dimer (ILID) for Controlling the Localization and Activity of Signaling Proteins. Proceedings of the National Academy of Sciences of the United States of America 2015, 112 (1), 112–117. https://doi.org/10.1073/pnas.1417910112.

(10) Nihongaki, Y.; Kawano, F.; Nakajima, T.; Sato, M. Photoactivatable CRISPR-Cas9 for Optogenetic Genome Editing. Nature Biotechnology 2015, 33 (7), 755–760. https://doi.org/10.1038/nbt.3245.

(11) Van Haren, J.; Charafeddine, R. A.; Ettinger, A.; Wang, H.; Hahn, K. M.; Wittmann, T. Local Control of Intracellular Microtubule Dynamics by EB1 Photodissociation. Nature Cell Biology 2018, 20 (3), 252–261. https://doi.org/10.1038/s41556-017-0028-5.

(12) Yumerefendi, H.; Lerner, A. M.; Zimmerman, S. P.; Hahn, K.; Bear, J. E.; Strahl, B. D.; Kuhlman, B. Light-Induced Nuclear Export Reveals Rapid Dynamics of Epigenetic Modifications. Nature Chemical Biology 2016, 12 (6), 399–401. https://doi.org/10.1038/nchembio.2068.

(13) Johnson, H. E.; Toettcher, J. E. Signaling Dynamics Control Cell Fate in the Early Drosophila Embryo. Developmental Cell 2019, 48 (3), 361–370.e3. https://doi.org/10.1016/j.devcel.2019.01.009.

(14) Li, Y.; Zhao, M.; Wei, D.; Zhang, J.; Ren, Y. Photocontrol of Itaconic Acid Synthesis in Escherichia Coli. ACS Synthetic Biology 2022, 11 (6), 2080–2088. https://doi.org/10.1021/acssynbio.2c00014.

(15) Koganezawa, Y.; Umetani, M.; Sato, M.; Wakamoto, Y. History-Dependent Physiological Adaptation to Lethal Genetic Modification under Antibiotic Exposure. eLife 2022, 11, 1–25. https://doi.org/10.7554/eLife.74486.

(16) Spencer, D. M.; Wandless, T. J.; Schreiber, S. L.; Crabtree, G. R. Controlling Signal Transduction with Synthetic Ligands. Science 1993, 262 (5136), 1019–1024. https://doi.org/10.1126/science.769436.

(17) Tyszkiewicz, A. B.; Muir, T. W. Activation of Protein Splicing with Light in Yeast. Nature Methods 2008, 5 (4), 303–305. https://doi.org/10.1038/nmeth.1189.

(18) Levskaya, A.; Weiner, O. D.; Lim, W. A.; Voigt, C. A. Spatiotemporal Control of Cell Signalling Using a Light-Switchable Protein Interaction. Nature 2009, 461 (7266), 997–1001. https://doi.org/10.1038/nature08446.

(19) Kennedy, M. J.; Hughes, R. M.; Peteya, L. A.; Schwartz, J. W.; Ehlers, M. D.; Tucker, C. L. Rapid Blue-Light-Mediated Induction of Protein Interactions in Living Cells. Nature Methods 2010, 7 (12), 973–975. https://doi.org/10.1038/nmeth.1524.

(20) Kawano, F.; Suzuki, H.; Furuya, A.; Sato, M. Engineered Pairs of Distinct Photoswitches for Optogenetic Control of Cellular Proteins. Nature Communications 2015. https://doi.org/10.1038/ncomms7256.

(21) Sheets, M. B.; Wong, W. W.; Dunlop, M. J. Light-Inducible Recombinases for Bacterial Optogenetics. ACS Synthetic Biology 2020, 9 (2), 227–235. https://doi.org/10.1021/acssynbio.9b00395.

(22) Dagliyan, O.; Krokhotin, A.; Ozkan-Dagliyan, I.; Deiters, A.; Der, C. J.; Hahn, K. M.; Dokholyan, N. V. Computational Design of Chemogenetic and Optogenetic Split Proteins. Nature Communications 2018, 9 (1). https://doi.org/10.1038/s41467-018-06531-4.

(23) Mahdavi, A.; Segall-Shapiro, T. H.; Kou, S.; Jindal, G. A.; Hoff, K. G.; Liu, S.; Chitsaz, M.; Ismagilov, R. F.; Silberg, J. J.; Tirrell, D. A. A Genetically Encoded and Gate for Cell-Targeted Metabolic Labeling of Proteins. Journal of the American Chemical Society 2013, 135 (8), 2979–2982. https://doi.org/10.1021/ja400448f.

(24) Nadler, D. C.; Morgan, S.; Flamholz, A.; Kortright, K. E.; Savage, D. F. Rapid Construction of Metabolite Biosensors Using Domain-Insertion Profiling. Nature Communications 2016, 7 (1). https://doi.org/10.1038/ncomms12266.

(25) Engler, C.; Kandzia, R.; Marillonnet, S. A One Pot, One Step, Precision Cloning Method with High Throughput Capability. PLoS ONE 2008, 3 (11), e3647. https://doi.org/10.1371/journal.pone.0003647.

(26) Nagy, A. Cre Recombinase: The Universal Reagent for Genome Tailoring. Genesis 2000, 26 (2), 99–109. https://doi.org/10.1002/(SICI)1526-968X(200002)26:2<99::AID-GENE1>3.0.CO;2-B.

(27) Meinke, G.; Bohm, A.; Hauber, J.; Pisabarro, M. T.; Buchholz, F. Cre Recombinase and Other Tyrosine Recombinases. Chemical Reviews 2016, 116 (20), 12785–12820. https://doi.org/10.1021/acs.chemrev.6b00077.

(28) Kawano, F.; Okazaki, R.; Yazawa, M.; Sato, M. A Photoactivatable Cre-LoxP Recombination System for Optogenetic Genome Engineering. Nature Chemical Biology 2016, 12 (12), 1059–1064. https://doi.org/10.1038/nchembio.2205.

(29) Sun, Z.; Liu, Q.; Qu, G.; Feng, Y.; Reetz, M. T. Utility of B-Factors in Protein Science: Interpreting Rigidity, Flexibility, and Internal Motion and Engineering Thermostability. Chemical Reviews 2019. https://doi.org/10.1021/acs.chemrev.8b00290.

(30) Leman, J. K.; Weitzner, B. D.; Lewis, S. M.; Adolf-Bryfogle, J.; Alam, N.; Alford, R. F.; Aprahamian, M.; Baker, D.; Barlow, K. A.; Barth, P.; Basanta, B.; Bender, B. J.; Blacklock, K.; Bonet, J.; Boyken, S. E.; Bradley, P.; Bystroff, C.; Conway, P.; Cooper, S.; Correia, B. E.; Coventry, B.; Das, R.; De Jong, R. M.; DiMaio, F.; Dsilva, L.; Dunbrack, R.; Ford, A. S.; Frenz, B.; Fu, D. Y.; Geniesse, C.; Goldschmidt, L.; Gowthaman, R.; Gray, J. J.; Gront, D.; Guffy, S.; Horowitz, S.; Huang, P. S.; Huber, T.; Jacobs, T. M.; Jeliazkov, J. R.; Johnson, D. K.; Kappel, K.; Karanicolas, J.; Khakzad, H.; Khar, K. R.; Khare, S. D.; Khatib, F.; Khramushin, A.; King, I. C.; Kleffner, R.; Koepnick, B.; Kortemme, T.; Kuenze, G.; Kuhlman, B.; Kuroda, D.; Labonte, J. W.; Lai, J. K.; Lapidoth, G.; Leaver-Fay, A.; Lindert, S.; Linsky, T.; London, N.; Lubin, J. H.; Lyskov, S.; Maguire, J.; Malmström, L.; Marcos, E.; Marcu, O.; Marze, N. A.; Meiler, J.; Moretti, R.; Mulligan, V. K.; Nerli, S.; Norn, C.; Ó’Conchúir, S.; Ollikainen, N.; Ovchinnikov, S.; Pacella, M. S.; Pan, X.; Park, H.; Pavlovicz, R. E.; Pethe, M.; Pierce, B. G.; Pilla, K. B.; Raveh, B.; Renfrew, P. D.; Burman, S. S. R.; Rubenstein, A.; Sauer, M. F.; Scheck, A.; Schief, W.; Schueler-Furman, O.; Sedan, Y.; Sevy, A. M.; Sgourakis, N. G.; Shi, L.; Siegel, J. B.; Silva, D. A.; Smith, S.; Song, Y.; Stein, A.; Szegedy, M.; Teets, F. D.; Thyme, S. B.; Wang, R. Y. R.; Watkins, A.; Zimmerman, L.; Bonneau, R. Macromolecular Modeling and Design in Rosetta: Recent Methods and Frameworks. Nature Methods 2020, 17 (7), 665–680. https://doi.org/10.1038/s41592-020-0848-2.

(31) Perrakis, A.; Sixma, T. K. AI Revolutions in Biology. EMBO reports 2021, 22 (11), 1–6. https://doi.org/10.15252/embr.202154046.

(32) Coyote-Maestas, W.; Nedrud, D.; Okorafor, S.; He, Y.; Schmidt, D. Targeted Insertional Mutagenesis Libraries for Deep Domain Insertion Profiling. Nucleic Acids Research 2020, 48 (2), 1–14. https://doi.org/10.1093/nar/gkz1110.

(33) Lee, T. S.; Krupa, R. A.; Zhang, F.; Hajimorad, M.; Holtz, W. J.; Prasad, N.; Lee, S. K.; Keasling, J. D. BglBrick Vectors and Datasheets: A Synthetic Biology Platform for Gene Expression. Journal of Biological Engineering 2011, 5, 12. https://doi.org/10.1186/1754-1611-5-12.

(34) Gerhardt, K. P.; Olson, E. J.; Castillo-Hair, S. M.; Hartsough, L. A.; Landry, B. P.; Ekness, F.; Yokoo, R.; Gomez, E. J.; Ramakrishnan, P.; Suh, J.; Savage, D. F.; Tabor, J. J. An Open-Hardware Platform for Optogenetics and Photobiology. Scientific Reports 2016, 6 (November), 1–13. https://doi.org/10.1038/srep35363.

(35) Bolger, A. M.; Lohse, M.; Usadel, B. Trimmomatic: A Flexible Trimmer for Illumina Sequence Data. Bioinformatics 2014, 30 (15), 2114–2120. https://doi.org/10.1093/bioinformatics/btu170.

(36) Salvatier, J.; Wiecki, T. V.; Fonnesbeck, C. Probabilistic Programming in Python Using PyMC3. PeerJ Computer Science 2016, 2016 (4), 1–24. https://doi.org/10.7717/peerj-cs.55.

