## Supplementary Information for "Comprehensive screening of a light-inducible split Cre recombinase with domain insertion profiling"

**Supplementary Tables**

**Table S1**. Plasmids used in this study.

| *Plasmid* | *Origin* | *Operon* | *Resistance* | *Reference* |
|---|---|---|---|---|
| *pUCKanR-Mu-BsaI* | pMB1 | R1R2(BsaI)-$P_{Cat}$-catR-R2R1(BsaI) | $Cm^R$ | [1] |
| *pATT-Dest-RFP* | pMB1 | $P_{lac}$-lacZ, $P_{lacUV5}$-mRFP1 | $Amp^R$ | This study, derived from [1] |
| *pATT-Dest-RFP-Cre* | pMB1 | cre (BsmBI flanked), $P_{lacUV5}$-mRFP1 | $Amp^R$ | This study |
| *pTKEI-Dest-loxP-sfGFP* | ColE1 | $P_{lac}$-lacZ, $P_{W7}$-loxP-term.-loxP-sfGFP | $Kan^R$ | This study, derived from [1] |
| *pATT-BsaI-pMag-nMagHigh1* | pMB1 | linker-pMag-nMagHigh1-linker (BsaI flanked) | $Amp^R$ | This study |
| *pBbS8c-Cre* | SC101 | $P_{bad}$-cre | $Cm^R$ | This study, derived from pBbS8c-mRFP1 [2] |

**Table S2**. DNA sequences used in this study.

| ***Gene*** | ***Sequence*** |
|---|---|
| *Cre* | gccacctctgatgaagtcaggaagaacctgatggacatgttcagggacaggcaggccttctctgaacacacctgg<br>aagatgctcctgtctgtgtgcagatcctgggctgcctggtgcaagctgaacaacaggaaatggttccctgctgaac<br>ctgaggatgtgagggactacctcctgtacctgcaagccagaggcctggctgtgaagaccatccaacagcacctggg<br>ccagctcaacatgctgcacaggagatctggcctgcctcgcccttctgactccaatgctgtgtccctggtgatgagga<br>gaatcagaaaggagaatgtggatgctggggagagagccaagcaggccctggcctttgaacgcactgactttgacc<br>aagtcagatccctgatggagaactctgacagatgccaggacatcaggaacctggccttcctgggcattgcctacaa<br>caccctgctgcgcattgccgaaattgccagaatcagagtgaaggacatctcccgcaccgatggtgggagaatgctg<br>atccacattggcaggaccaagaccctggtgtccacagctggtgtggagaaggccctgtccctgggggttaccaagct<br>ggtggagagatggatctctgtgtctggtgtggctgatgaccccaacaactacctgttctgccgggtcagaaagaatg<br>gtgtggctgccccttctgccacctcccaactgtccacccgggccctggaagggatctttgaggccacccaccgcctga<br>tctatggtgccaaggatgactctgggcagagatacctggcctggtctggccactctgccagagtgggtgctgccagg<br>gacatggccagggctggtgtgtccatccctgaaatcatgcaggctggtggctggaccaatgtgaacattgtgatgaac<br>tacatcagaaacctggactctgagactggggccatggtgaggctgctcgaggatggggac |
| *Magnet insert(linker-pMag-RBS-nMagHigh1-linker)* | gcatcgggttctggaggctcaagcggatcacacactctttacgcccctggaggatacgacattatgggatatt<br>tgcggcagattaggaaccgcccaaaccctcaggtcgaactggggcctgtggacacgtcatgtgccctgatcctgtgcg<br>atctgaagcaaaaggacactccgatcgtctacgcctcggaagccttcttgtatatgaccggatacagcaatgcagagg<br>tgctcggcaggaactgcagattcctgcagtcccccgacgggatggtgaaaccaaagtcgactcgcaaatatgtggact<br>cgaacacgatcaacaccatgcggaaggccatcgaccggaacgccgaggtccaggtggaggtggtcaactttaagaag<br>aacggccagcggttcgtgaactttctgaccatgattccggtccgggatgaaaccggagagtacagatactccatgggat<br>tccagtgcgaaacagaataagaattcattaaagaggagaaaggtaccatgcatacactttacgctcctgggggctacga<br>catcatgggctatttggatcagattggcaatcgcccgaatccacaggttgaattagggccagtcgatacgtcgtgcgcact<br>gattttgtgtgatttaaagcaaaaggataccccaattgtttacgcgagtgaggcgtttctgtatatgacgggctactcaaat<br>gcggaggtacttggccgcaactgtcgcttcttacaatcgccggacggcatggtaaagcctaagtcaactcgtaaatacgt<br>tgactccaacactatcaatacaattcgcaaagcgatcgatcgcaacgcagaggtccaggtggaggttgttaactttaaga<br>agaatgggcaacgcttcgtgaattttcttacgattattccggttcgtgacgaaaccggcgaatatcgttactctatggggtt<br>ccagtgtgaaaccgaaggtggcggaggtagcgcgt |
| *Cre reporter ($P_{W7}$-loxP-terminator-loxP-sfGFP)* | ttatcaaaaagagtattgaaataaagtctaacctataggaagattacagccatcgagagggacacggcgaa<br>ataacttcgtatagcatacattatacgaagttatccaggcatcaaataaggatccaaactcgagtaaggatctccag<br>gcatcaaataaaacgaaaggctcagtcgaaagactgggcctttcgttttatctgttgtttgtcggtgaacgctctctac<br>tagagtcacactggctcaccttcgggtgggcctttctgcgtttataccataacttcgtatagcatacattatacgaagtt<br>attttaagaaggagatatacatatgcgtaaaggcgaagagctgttcactggtgtcgtccctattctggtggaact<br>ggatggtgatgtcaacggtcataagttttccgtgcgtggcgagggtgaaggtgacgcaactaatggtaaactgacgct<br>gaagttcatctgtactactggtaaactgccggtaccttggccgactctggtaacgacgctgacttatggtgttcagtgct<br>ttgctcgttatccggaccatatgaagcagcatgacttcttcaagtccgccatgccggaaggctatgtgcaggaacgcac<br>gatttcctttaaggatgacggcacgtacaaaacgcgtgcggaagtgaaatttgaaggcgataccctggtaaaccgcat<br>tgagctgaaaggcattgactttaaagaagacggcaatatcctgggccataagctggaatacaattttaacagccacaa<br>tgtttacatcaccgccgataaacaaaaaaatggcattaaagcgaattttaaaattcgccacaacgtggaggatggcag<br>cgtgcagctggctgatcactaccagcaaaacactccaatcggtgatggtcctgttctgctgccagacaatcactatctg<br>agcacgcaaagcgttctgtctaaagatccgaacgagaaacgcgatcatatggttctgctggagttcgtaaccgcagc<br>gggcatcacgcatggtatggatgaactgtacaaataa |

| | |
|---|---|
| *Modified transposon* | tgcatcggagaccgaaaaacgcgaaagcgtttcacgataaatgcgaaaacggatcgatccttttcgaccgaataaata<br>cctgtgacggaagatcacttcgcagaataaataaatcctggtgtccctgttgataccgggaagccctgggccaacttttg<br>gcgaaaatgagatgttgatcggcacgtaagaggttccaactttcaccataatgaaataagatcactaccgggcgtatttt<br>ttgagttgtcgagattttcaggagctaaggaagctaaaatggagaaaaaaatcactggatataccaccgttgatatatcc<br>caatggcatcgtaaagaacattttgaggcatttcagtcagttgctcaatgtacctataaccagaccgttcagctggatatt<br>acggcctttttaaagaccgtaaagaaaaataagcacaagttttatccggcctttattcacattcttgcccgcctgatgaat<br>gctcatccggaattacgtatggcaatgaaagacggtgagctggtgatatgggatagtgttcacccttgttacaccgttttcc<br>atgagcaaactgaaacgttttcatcgctctggagtgaataccacgacgatttccggcagtttctacacatatattcgcaaga<br>tgtggcgtgttacggtgaaaacctggcctatttccctaaagggtttattgagaatatgtttttcgtgtcagccaatccctgggt<br>gagtttcaccagttttgatttaaacgtggccaatatggacaacttcttcgcccccgttttcaccatgggcaaatattatacgca<br>aggcgacaaggtgctgatgccgctggcgattcaggttcatcatgccgtttgtgatggcttccatgtcggcagaatgcttaatg<br>aattacaacagtactgcgatgagtggcagggcggggcgtaatttttttaaggcagttattggtgcccttaaacgcctggttgc<br>tacgcctgaataagtgataataagcggatgaatggcagaaattcgaaagcaaattcgacccggtcgtcggttcagggcag<br>ggtcgttaaatagccgcttatgtctattgctggtttaccggtttattgactaccggaagcagtgtgaccgtgtgcttctcaaatg<br>cctgaggccagtttgctcaggctctccccgtggaggtaataattgacgataggatcgatccgttttcgcatttatcgtgaaacg<br>ctttcgcgtttttcggtctccgcgtca |

**Table S3**. Amino acid sequences. * Indicates stop codon. (RBS) indicates a ribosome binding site

| ***Protein*** | ***Sequence*** |
|---|---|
| *Cre (indexed 1-326)* | ATSDEVRKNLMDMFRDRQAFSEHTWKMLLSVCRSWAAWCKLNNRKWFPAEPEDVRD<br>YLLYLQARGLAVKTIQQHLGQLNMLHRRSGLPRPSDSNAVSLVMRRIRKENVDAGERAK<br>QALAFERTDFDQVRSLMENSDRCQDIRNLAFLGIAYNTLLRIAEIARIRVKDISRTDGGRML<br>IHIGRTKTLVSTAGVEKALSLGVTKLVERWISVSGVADDPNNYLFCRVRKNGVAAPSATSQ<br>LSTRALEGIFEATHRLIYGAKDDSGQRYLAWSGHSARVGAARDMARAGVSIPEIMQAGG<br>WTNVNIVMNYIRNLDSETGAMVRLLEDGD |
| *Magnet insert* (*linker*-pMag-*RBS*-nMagHigh1-*linker*) | ASGSGGSSGSHTLYAPGGYDIMGYLRQIRNRPNPQVELGPVDTSCALILCDLKQKDTPIVY<br>ASEAFLYMTGYSNAEVLGRNCRFLQSPDGMVKPKSTRKYVDSNTINTMRKAIDRNAEVQ<br>VEVVNFKKNGQRFVNFLTMIPVRDETGEYRYSMGFQCETE * *(RBS)* MHTLYAPGGY<br>DIMGYLDQIGNRPNPQVELGPVDTSCALILCDLKQKDTPIVYASEAFLYMTGYSNAEVLGR<br>NCRFLQSPDGMVKPKSTRKYVDSNTINTIRKAIDRNAEVQVEVVNFKKNGQRFVNFLTIIP<br>VRDETGEYRYSMGFQCETEGGGGSAS |

### Supplementary Figures

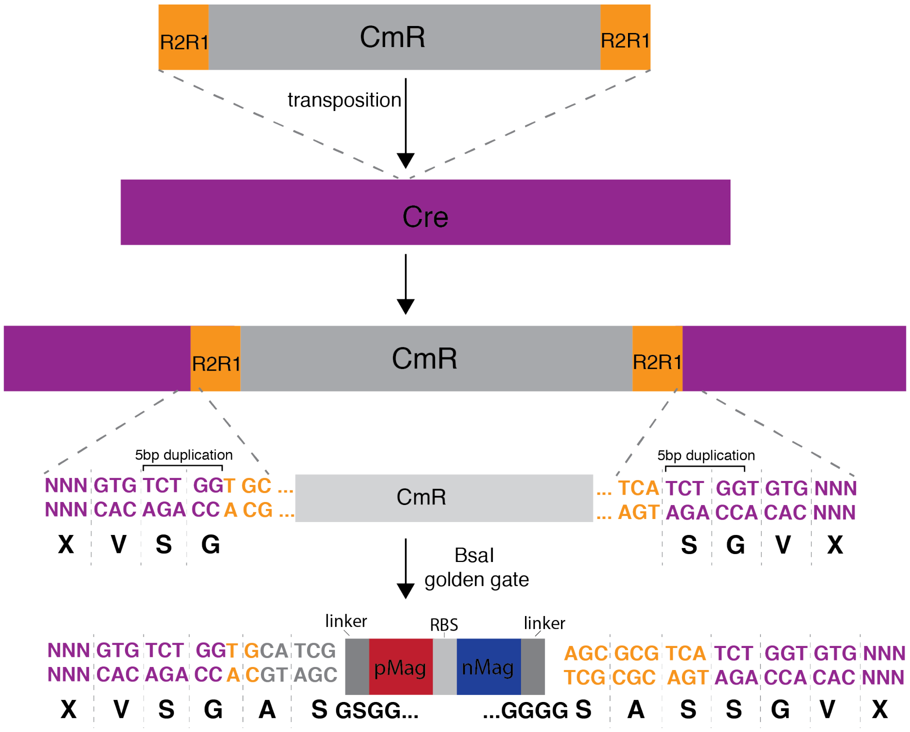


**Figure S1**. Schematic of transposon insertion. MuA-mediated transposition results in a 5 nucleotide duplication of the Cre sequence and addition of a 2 nucleotide scar on the 5' and 3' ends of the transposon. Replacement of the transposon with the magnet dimer domains through a BsaI golden gate maintains the open reading frame with flexible linkers connecting N- and C-terminal fragments of Cre with pMag and nMagHigh1, respectively. R2R1 sites are sequences recognized by MuA transposase and necessary for transposition.

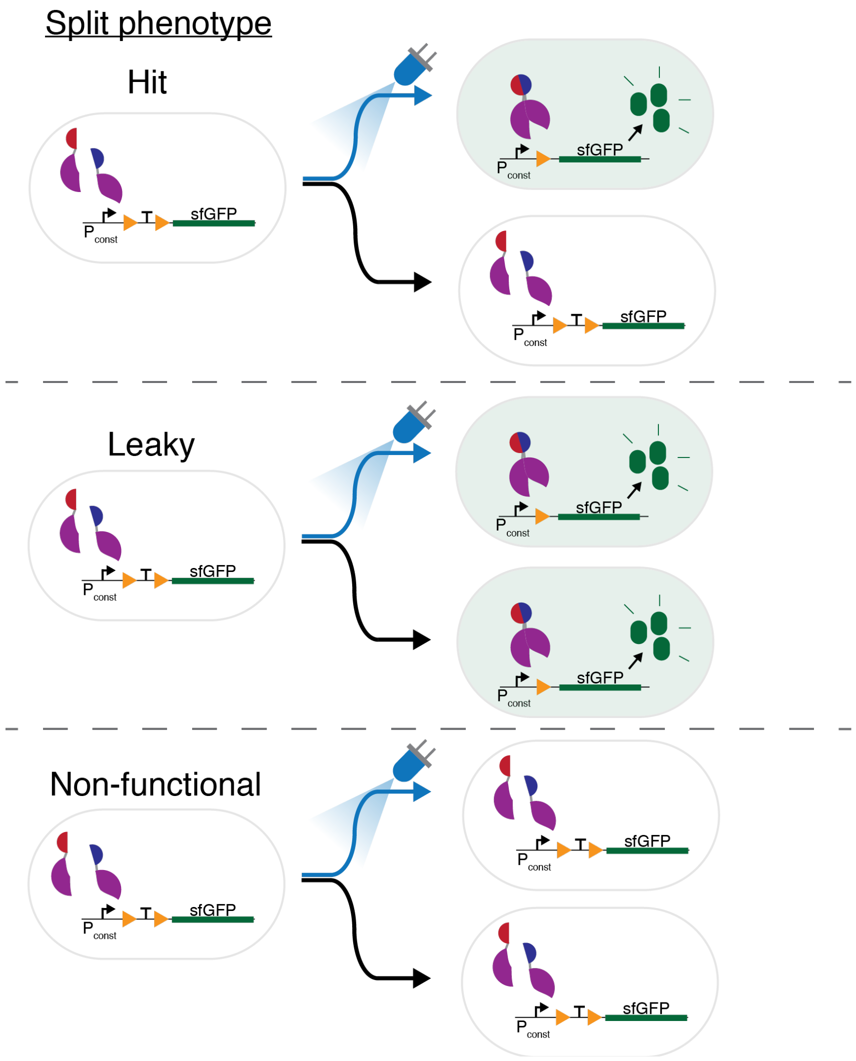


**Figure S2**. A split Cre construct can function as one of three phenotypes using the sfGFP Cre reporter: 1) Hit: Cre recombinase activity is light responsive. 2) Leaky: Cre is active even in the dark, 3) Non-functional: Cre is inactive in dark and light conditions.

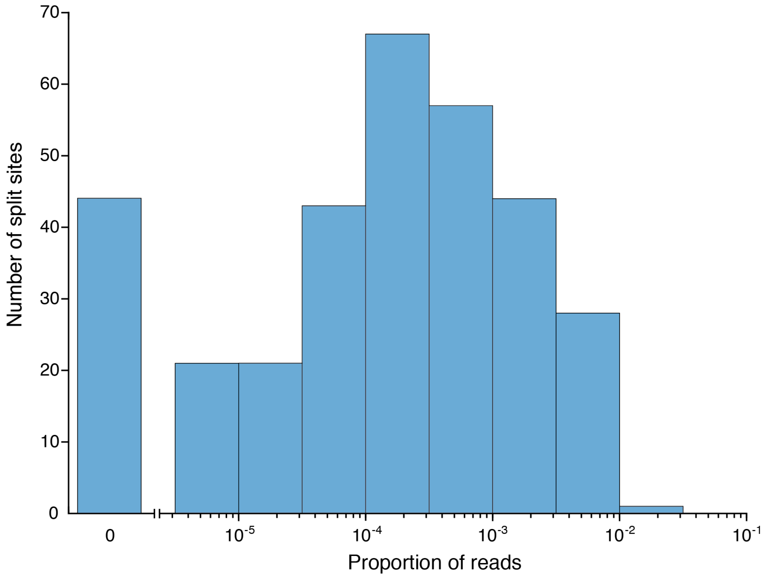


**Figure S3**. Distribution of split sites in the light-exposed naïve library prior to fluorescent activated cell sorting.

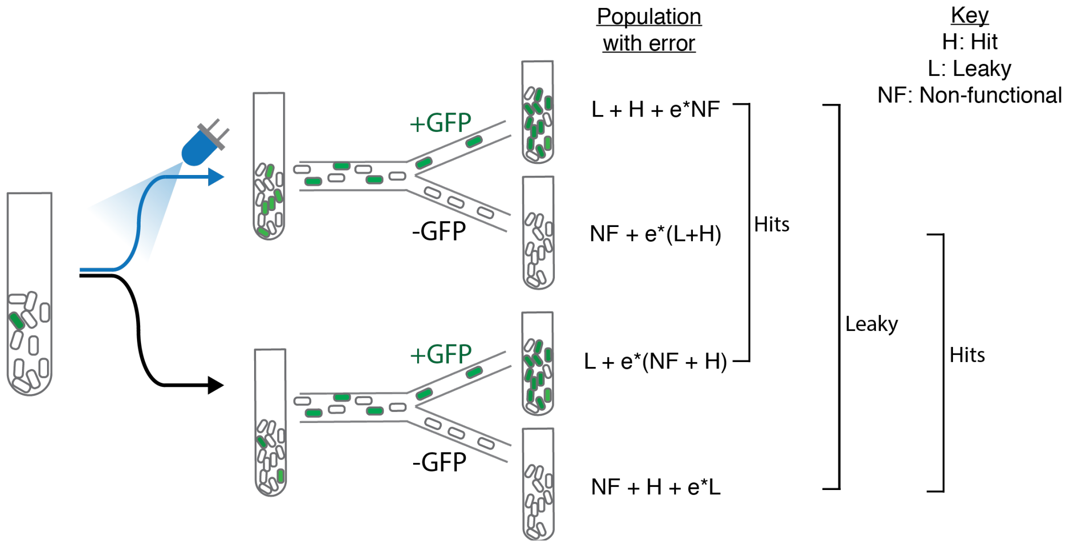


**Figure S4**. Schematic detailing the makeup of split phenotypes expected to be present in each population after sorting. Sorting error, e, leads to spurious events, e.g. non-functional splits present in the +GFP populations. Three ratio comparisons of the resulting distributions can be used to extract information from the libraries to overcome variation stemming from sorting errors and under sampled variants. Light/+GFP compared to Dark/+GFP will reveal hits, Light/+GFP compared to Dark/-GFP provides information about possible leaky variants, and Light/-GFP compared to Dark/-GFP will reveal hits.

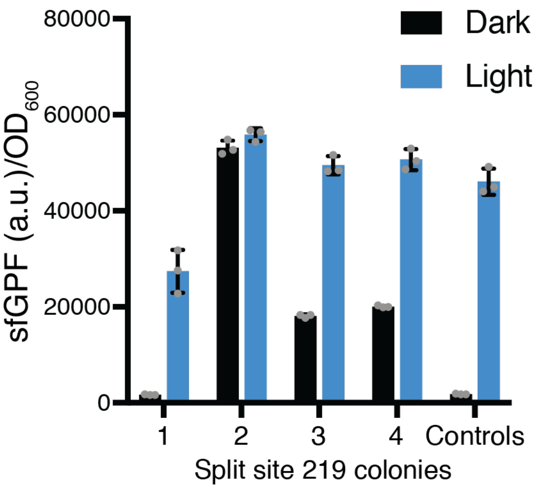


**Figure S5**. sfGFP expression levels with and without exposure to 465 nm blue light in four colonies of split Cre 219 tested in parallel. Controls display sfGFP expression from the reporter with and without the addition of functional, non-split Cre. Error bars show standard deviation around the mean (n = 3 replicates).

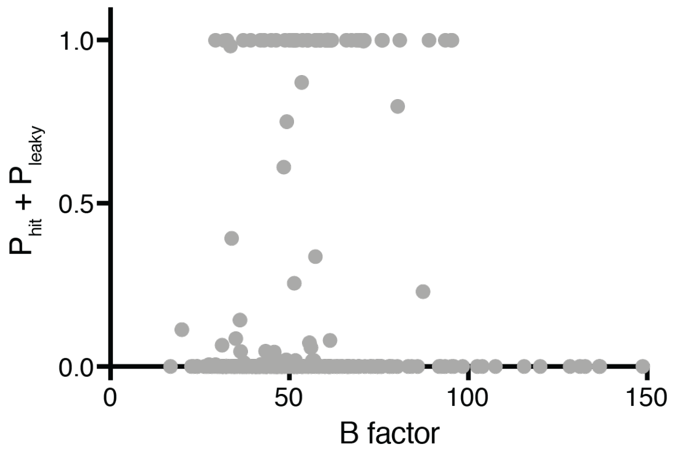


**Figure S6**. Probability of catalytic activity (hit or leaky) plotted against B-factor for residues in the Cre structure (PDB: 1NZB).

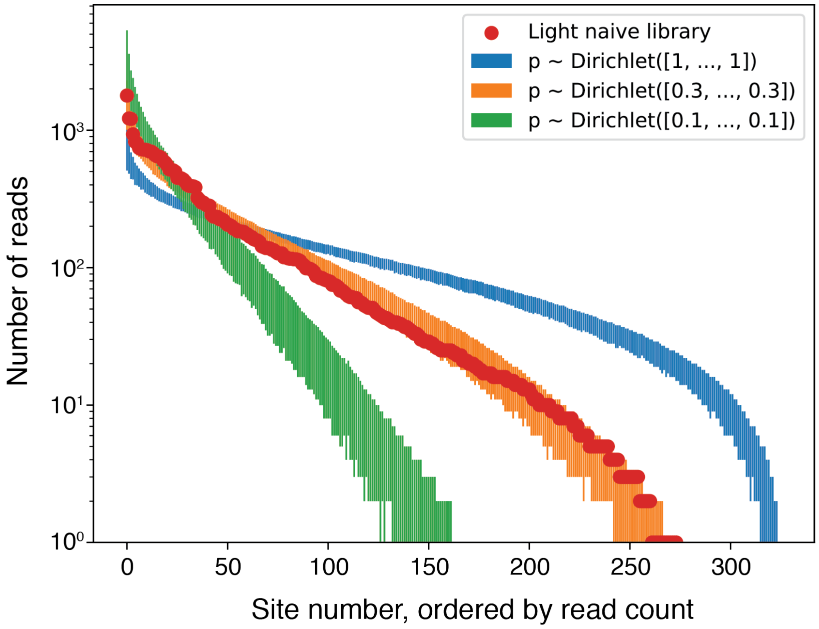


**Figure S7.** Choosing a Dirichlet prior of parameter [0.3, … , 0.3] for the naïve library distribution reproduces the experimentally observed dispersion well. The bars represent the 94% highest-density interval of the simulated results.

**Supplemental Text – Bayesian Modeling**

*Bayesian modeling*

We developed the statistical model as a generative model, i.e. a mapping in probabilistic terms of our knowledge of the experimental process. The experiment is highly random, and every step creates a probability distribution related to some of the previous distributions. The model creates the distributions and describes how they relate to each other.

*Nature of the split sites*

Every site $i \in [1, 326]$ can be a hit (H), leaky (L), or non-functional (N) variant. This is described by a categorical distribution. We give prior probabilities to H, L, and N according to the proportions: 2% hits, 8% leaky, and 90% non-functional, which is based on empirical observations.

$$site_i \sim \text{Categorical}(H : 2\%, L : 8\%, N : 90\%)$$

*Distribution of the naïve populations*

The split sites are generated by a random process with a non-uniform distribution. We adapted the Dirichlet probability distribution, to describe a prior over this initial split site distribution. The parameter of a Dirichlet distribution is a vector of N numbers, where N is the number of categories (here 326 possible split sites). Because we do not want to favor any split site a priori, we leave all 326 numbers equal to the same value, the only adjustment being the shared value $\alpha$. We create two naïve distributions: $p_L$ for the light naïve library, and $p_D$ for the dark naïve library.

$$p_{L,D} \sim \text{Dirichlet}\ ([\alpha, \dots, \alpha])$$

Modeling the plasmid sequencing process as a multinomial sampling over the naïve library, we can describe the naïve light and dark read counts as follows, where $N$ is the total number of read counts (sequencing depth):

$$(L, D) \sim \text{Multinomial}\ (N, p_{L,D})$$

Where L and D are vectors of 326 length with read counts corresponding to the abundance of a given variant. This allows us to simulate read counts that we would observe from Dirichlet-distributed naïve distributions (Fig. S7). Comparing these simulations to experimental data allowed us to determine an appropriate value for $\alpha$, where $\alpha$ = 0.3 reproduces the dispersion observed experimentally in the sequencing of the naïve libraries well.

*Sorting process*

The sorting process happens independently for every cell that passes through the cell sorter.

Light-treated cells that behave as hits (H) or leaky (L) are sorted as +GFP with a probability $1 - fn$, and -GFP with a probability $fn$, where $fn$ is the false negative rate. Inversely, light-treated cells that behave as non-functional (N) are sorted as +GFP with a probability $fp$, and -GFP with a probability $1 - fp$, where $fp$ is the false positive rate. We fixed both $fn$ and $fp$ at 25%, following empirical sorting observations.

Introducing $(s_L^{+,-})_i$ the probability that a split site at position $i$ and light-treated gets sorted as +,-GFP, we can write

$$(s_L^+)_i = \begin{cases} 1 - \text{fn} & \text{if sites}_i = L \text{ or } H \\ \text{fp} & \text{if sites}_i = N \end{cases}$$

and

$$(s_L^-)_i = 1 - (s_L^+)_i$$

Similarly, for the dark-treated population,

$$(s_D^+)_i = \begin{cases} 1 - \text{fn} & \text{if sites}_i = L \\ \text{fp} & \text{if sites}_i = N \text{ or } H \end{cases}$$

and

$$(s_D^-)_i = 1 - (s_D^+)_i$$

From there, we derive the proportions of each cell type in the four sorted populations. For split site $i$:

$$(p_{L,D}^{+,-})_i = \frac{(p_{L,D})_i (s_{L,D}^{+,-})_i}{\sum_j (p_{L,D})_j (s_{L,D}^{+,-})_j}$$

This allows us to finally express the sequencing results of the four sorted populations:

$$(L, D)^{+,-} \sim \text{ Multinomial } (N, p_{L,D}^{+,-})$$

*Model implementation*

We implemented the model in Python, with the PyMC library. The crux of the implementation is as follows:

```
with pm.Model():
    # 0 = H, 1 = L, 2 = N
    sites = pm.Categorical("sites", [p_hit, p_leaky, 1-p_hit-p_leaky], size=326)
    p_l = pm.Dirichlet("p_l", [alpha] * 326)
```

```
    p_d = pm.Dirichlet("p_d", [alpha] * 326)
    pm.Multinomial("L", n=sum(df.light), observed=df.light)
    pm.Multinomial("D", n=sum(df.dark), observed=df.dark)
    s_l = pm.math.stack([1 - fn_rate, 1 - fn_rate, fp_rate])[sites]
    pm.Multinomial("L+", n=sum(df.light_pos),
        p=p_l * s_l / (p_l * s_l).sum(), observed=df.light_pos)
    pm.Multinomial("L-", n=sum(df.light_neg),
        p=p_l * (1 - s_l) / (p_l * (1 - s_l)).sum(), observed=df.light_neg)

s_d = pm.math.stack([fp_rate, 1 - fn_rate, fp_rate])[sites]
pm.Multinomial("D+", n=sum(df.dark_pos),
    p=p_d * s_d / (p_d * s_d).sum(), observed=df.dark_pos)
pm.Multinomial("D-", n=sum(df.dark_neg),
    p=p_d * (1 - s_d) / (p_d * (1 - s_d)).sum(), observed=df.dark_neg)
```

*Inference*

We ran 4 Markov chain Monte Carlo simulations of 10,000 samples each, preceded by a phase of burn-in/tuning of the same length.

**References**

(1) Nadler, D. C.; Morgan, S.; Flamholz, A.; Kortright, K. E.; Savage, D. F. Rapid Construction of Metabolite Biosensors Using Domain-Insertion Profiling. *Nat. Commun.* **2016**, *7* (1). https://doi.org/10.1038/ncomms12266.

(2) Lee, T. S.; Krupa, R. A.; Zhang, F.; Hajimorad, M.; Holtz, W. J.; Prasad, N.; Lee, S. K.; Keasling, J. D. BglBrick Vectors and Datasheets: A Synthetic Biology Platform for Gene Expression. *J. Biol. Eng.* **2011**, *5*, 12. https://doi.org/10.1186/1754-1611-5-12.
